# PDE-constrained optimization for estimating population dynamics over cell cycle from static single cell measurements

**DOI:** 10.1101/2020.03.30.015909

**Authors:** Karsten Kuritz, Alain R Bonny, João Pedro Fonseca, Frank Allgöwer

**Affiliations:** Institute for Systems Theory and Automatic Control, University of Stuttgart, Stuttgart, 70569, Germany; Department of Biochemistry and Biophysics, California Institute for Quantitative Biosciences, University of California, San Francisco, CA 94158, USA; Chan Zuckerberg Biohub, San Francisco, CA 94158, USA; Stuttgart Research Center Systems Biology, University of Stuttgart, Stuttgart, 70569, Germany

## Abstract

**Motivation:** Understanding how cell cycle responds and adapts dynamically to a broad range of stresses and changes in the cellular environment is crucial for the treatment of various pathologies, including cancer. However, measuring changes in cell cycle progression is experimentally challenging, and model inference computationally expensive.

**Results:** Here, we introduce a computational framework that allows the inference of changes in cell cycle progression from static single-cell measurements. We modeled population dynamics with partial differential equations (PDE), and derive parameter gradients to estimate time- and cell cycle position-dependent progression changes efficiently. Additionally, we show that computing parameter sensitivities for the optimization problem by solving a system of PDEs is computationally feasible and allows efficient and exact estimation of parameters. We showcase our framework by estimating the changes in cell cycle progression in K562 cells treated with Nocodazole and identify an arrest in M-phase transition that matches the expected behavior of microtubule polymerization inhibition.

**Conclusions:** Our results have two major implications: First, this framework can be scaled to high-throughput compound screens, providing a fast, stable, and efficient protocol to generate new insights into changes in cell cycle progression. Second, knowledge of the cell cycle stage- and time-dependent progression function allows transformation from pseudotime to real-time thereby enabling real-time analysis of molecular rates in response to treatments.

**Availability:** MAPiT toolbox (Karsten Kuritz 2020) is available at github: https://github.com/karstenkuritz/MAPiT.

## 1 Introduction

Cells are regularly exposed to various stresses, including environmental changes or drug treatment. One of the first cell responses to environmental stresses is to modulate its cell cycle, usually stopping its progression. Cell cycle arrest occurs at any of the four cell cycle stages (G1, S, G2, M), and it is dynamically regulated (Kapuy et al. 2009).

Identifying cell cycle progression changes caused by environmental factors, or treatments, is crucial for the understanding of cellular responses and the development of new treatment options for many pathologies. Several algorithms designed to reconstruct cell trajectories from single-cell data, including Wanderlust, Monocle and diffusion maps (DPT), have been recently proposed (Saelens et al. 2019; Bendall et al. 2014; Theis et al. 2015; Qiu et al. 2017). These methods order single-cell data in pseudotime - a quantitative measure of progress through a biological process. We recently developed MAPiT, a computational framework that transforms pseudotime into real-time (Karsten Kuritz, Stöhr, et al. 2020; Karsten Kuritz 2020).

Here we apply MAPiT to detect dynamically-regulated changes in cell cycle progression by learning a time- and cell cycle position-dependent function from experimentally observed cell distributions (Figure 1). First, we introduce the experimental data and data processing steps, which provide the basis for our estimation algorithm. Then, we present the partial differential equation (PDE) to model cell cycle-dependent cell density and the cost function that form the PDE-constrained optimization problem. Our newly derived sensitivity system enables efficient solving of the problem. Finally, we demonstrate the capability of the algorithm with one artificial and two experimental datasets.

**Figure 1:**
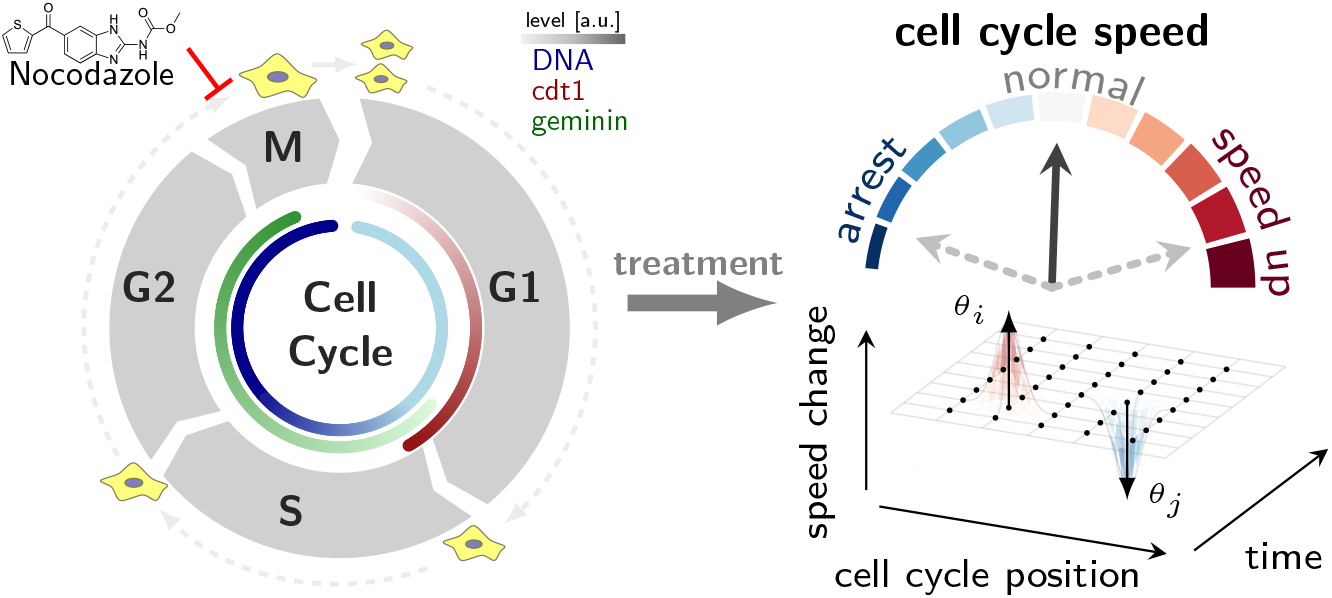
Schematic cell cycle progression with markers (DNA, cdt1, geminin) and Nocodazole treatment. A treatment causes time- and cell cycle position-dependent change in the cell cycle speed.

## 2 Results

### 2.1 Preprocessing of experimental data

To enable the identification of changes in cell cycle progression, we preprocessed the static data from time series single-cell experiments (Figure 2). In summary, we (1) found the path through the dataspace using pseudotime algorithms; (2) derived cell density on the path; (3) transformed to cell cycle position-dependent cell density with MAPiT; and (4) calculated the relative cell number density.

**Figure 2:**
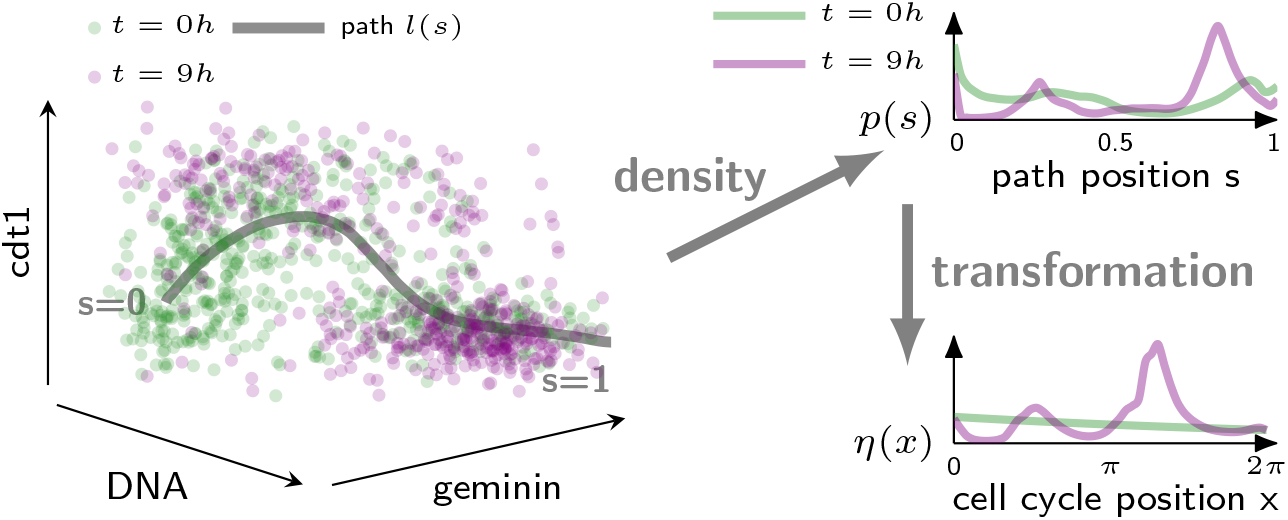
Data preprocessing steps shown for 0 and 9 *h* treatment with Nocodazole: Pseudotime algorithms find a path in the dataspace *l*(*s*) which an average cells takes during cell cycle progression. Kernel density estimation followed by dimensionality reduction provides a cell density on the path *p*(*s*). Applying a transformation *τ*(*s*) learned from unperturbed populations results in cell densities on a cell cycle position scale *η*(*x*).

To find the path in the dataset at *t* = 0 *h*, we applied a previously described pseudotime algorithm (Bendall et al. 2014). We probed different root cells for the pseudotime axis, and created the average path that a cell takes through the dataspace *l*: [0,1] → ℝ^3^ by a moving average of signal intensities along the pseudotime axis. To obtain the cell density *p*(*s*) along the path, we constructed multi-dimensional Gaussian distributions centered on the data coordinates *μ_i_* of each cell

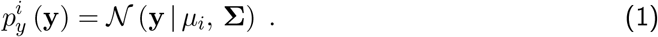

Bandwidths Σ were set with Silverman’s rule of thumb (Silverman 1986). We evaluated these distributions for each cell on the path *l*(*s*) and normalized resulting densities to 1.

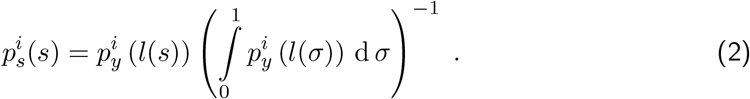

The cell number density along the path was given by the sum of all individual probability densities normalized by the number of cells.

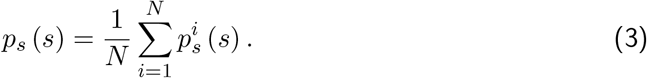

The density on the path *p_s_*(*s*) depends on various factors, such as the measured markers or the cell type. A transformation from the path position scale to an independent cell cycle position scale (normalized time scale) solves this issue. Applying MAPiT, we derived this transformation *τ*: *s* → *x* with 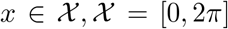. The densities at 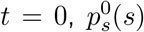, were transformed to the steady-state distribution of an unperturbed population of cycling cells (Powell 1956; Kafri et al. 2013; K. Kuritz et al. 2017; Karsten Kuritz, Stöhr, et al. 2020)

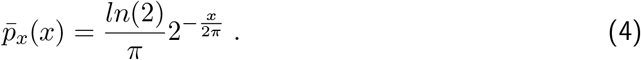

The cumulative density functions of the distributions were used to obtain the transformation

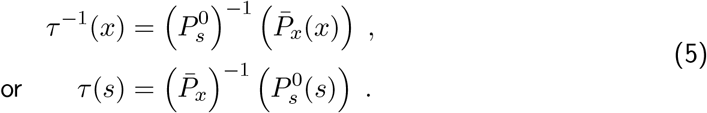

Applying this transformation to the density on the path at every time point, we uncover the distribution of cells in the cell cycle.

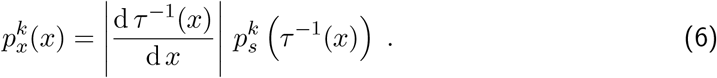

The procedure is illustrated for two time points in Figure 2 for a three-dimensional dataspace. Finally, we derived the cell number density at time points *t_k_* by multiplication of the densities 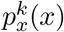 with the respective population size *N_k_*:

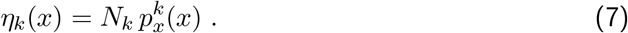

### 2.2 Model and optimization problem

We then asked what are the time- and cell cycle position-dependent changes in cell cycle progression speed, that reproduce experimentally observed time-course of cell number densities. To find the speed change function, we ran an optimization protocol that minimized the difference between the cell number densities from experimental data *η_k_*(*x*) and the model prediction *n_k_*(*x*). The optimization was constrained by requiring that the predicted number density is a solution of the age-structured population model type PDE

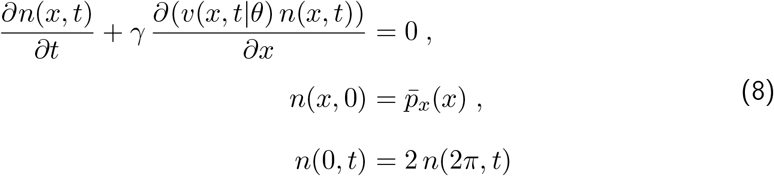

with 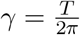, where *T* is the period of the cell cycle.

The speed change function *ν*(*x,t|θ*) is such that *ν*(*x, 0|θ*) = 1 which resulted in the classical age-structured population model for *t* = 0*h* (K. Kuritz et al. 2017; Foerster 1959). Parameters *θ* ∈ ℝ^*m*^ are the weights of *m* 2D normal distributions centered at grid points (*μ_i_*)_*i*=1,.,.,*m*_ in the *t* and *x* domains. To ensure the condition for *t* = 0 *h,* we defined the speed change function as the exponential of a product of a sum of Gaussians and a hill function in *t*:

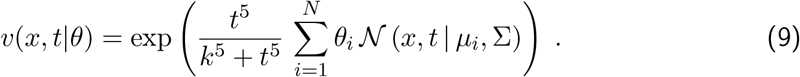

The bandwidth was chosen manually, depending on the number and spacing of grid points. An example of a speed change function is shown in Figure 1.

We used symmetric Kullback-Leibler divergence 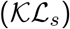 to quantify the deviation of model predictions from experimental data. The objective function for the parameter estimation was set as the sum of the logarithm of the 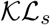 at all timepoints. Suppose, *p*(*x*) and *q*(*x*) are two probability distributions; then the symmetric Kullback-Leibler divergence is given by

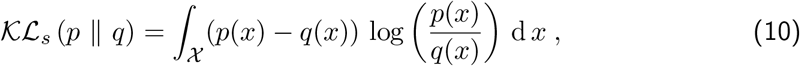

then the objective function reads

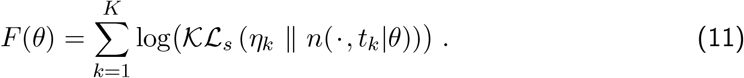

To minimize the difference between data *η_k_*(*x*) and model prediction *n*(*x,t_k_|θ*), we pose the following optimization problem:

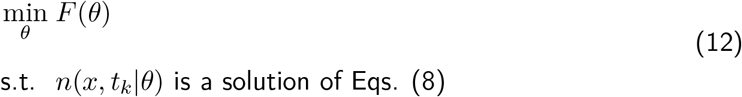

This PDE-constrained optimization problem is non-convex and possesses several local min-ima.

### 2.3 Sensitivity system for efficient parameter estimation

The availability of gradients for the objective function concerning the parameters dramatically improves the solution of optimization problems. These gradients can be approximated by finite difference, or computed as the solution of a sensitivity system. Derivative-based optimization employing the sensitivity equations is known to outperform other methods by orders of magnitude in accuracy and speed (Raue et al. 2013). To enable the efficient parameter estimation for the PDE-constrained optimization problem, we showed the gradient of the objective function

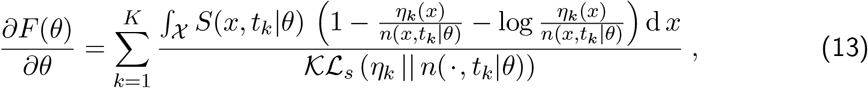

with 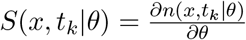 denoting the sensitivity of the predicted number density at time point *t_k_* for parameters *θ*.

To increase clarity in the derivation of the sensitivity system, we used a shorthand notation where we omitted the arguments in the functions *S,n,v*. Differentiating the system in Eqs. (8) with respect to the parameters *θ* provided a PDE for the sensitivities

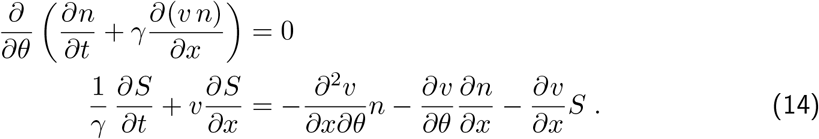

In classical ODE constrained optimization problems, the sensitivities are obtained by solving a system of differential equations (Raue et al. 2013). Unlike in ODE sensitivity systems, closure of the differential equations was not achieved because the unknown function 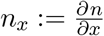 emerged. Differentiating the system in Eqs. (8) with respect to *x* results in a new PDE:

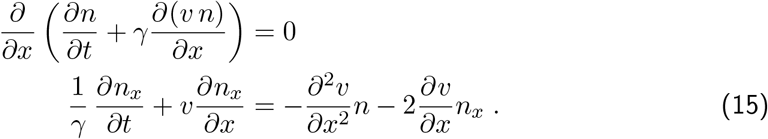

The system of coupled first order nonhomogenous PDEs composed of Eqs. (8),(14),(15) was used to simulate the sensitivities which we applied in Eq. (13) to compute the gradient of the objective function. The resulting system of PDEs is given by:

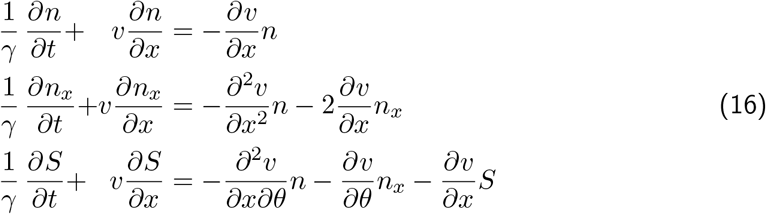

With initial and boundary conditions

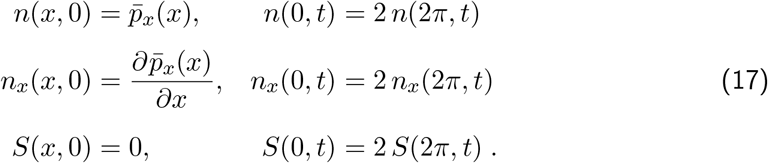

### 2.4 Practical realization

Two concepts are critical for the implementation of computational methods. First, they rely on assumptions about the biological system and experimental setup. Second, they must be computationally feasible (i.e. do not need a large amount of time to generate results).

The following assumptions underlie the algorithm as presented here:

**Assumption 1** *Cell death and cell cycle arrest cannot be distinguished unambiguously. For example, the same cell densities can result from all cells slowing down or just a few cells dying.*
**Assumption 2** *Treatment only changes velocity on path, not direction/route of path in dataspace.*
**Assumption 3** *Response to the stress at specific cell cycle stage is homogeneous.*

We verified assumptions 1 and 2 by excluding dead cells, using the light scattering characteristics of live cells and by manual inspection of data and path as shown in Figure 2.

Solving large PDE systems is computationally expensive. However, by discretizing the solution in *x* in *n* points, we transfered the problem to *n* decoupled ODE systems by the method of characteristics (Evans 2010).

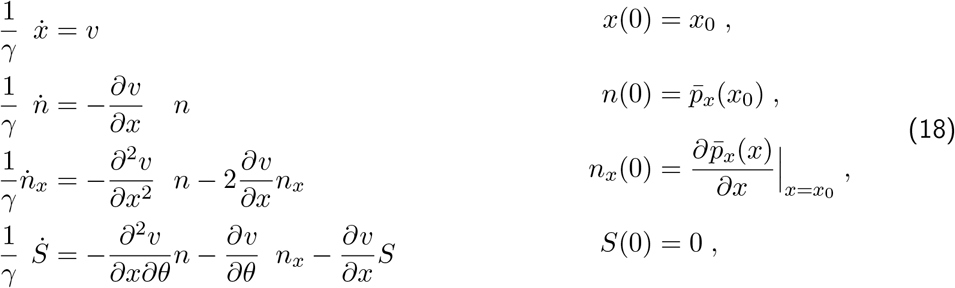

The linear time varying (LTV) ODE system in Eqs. (18) had discontinuities at division events where *x*(*t*) = 2*π*. We used the AMICI toolbox for simulation of system Eqs. (18) (Fröhlich et al. 2017) because it is optimized for efficiently solving large systems of ODE. The LTV system Eqs. (18) has for a single discretization, 3 + *m* states, with *m* denoting the number of parameters. Thus, computational complexity increases as a product of the number of parameters, discretization points, and time points 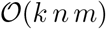. In the examples with experimental data we have *k* = 6,*n* = 50, *m* = 64. A Single evaluation of the objective function in Matlab R2018b on a laptop with Linux, processor Intel Core i5-4300U CPU @ 1.9GHz x 4, 8 GB RAM, took 3.5 or 0.5 seconds, with or without sensitivities, respectively.

### 2.5 Evaluation with in silico data

We applied our method to recover parameters from an artificial speed change function. We generated a simple speed change function composed of *m* = 9 2D Gaussian distributions with arbitrarily chosen weights (Figure 3). Forward simulation of model Eqs. (8) provided the cell number densities of an in silico experiment. Estimating the parameters resulted in an almost perfect fit of the predictions of the fitted model with the in silico-generated data. Furthermore, estimated parameter values, and the resulting speed change function strongly resembled those underlying the in silico data (Figure 3). This example demonstrated the capability of the PDE-constrained estimation approach to correctly estimate an unknown speed change function from a time course experiment with cell number densities.

**Figure 3:**
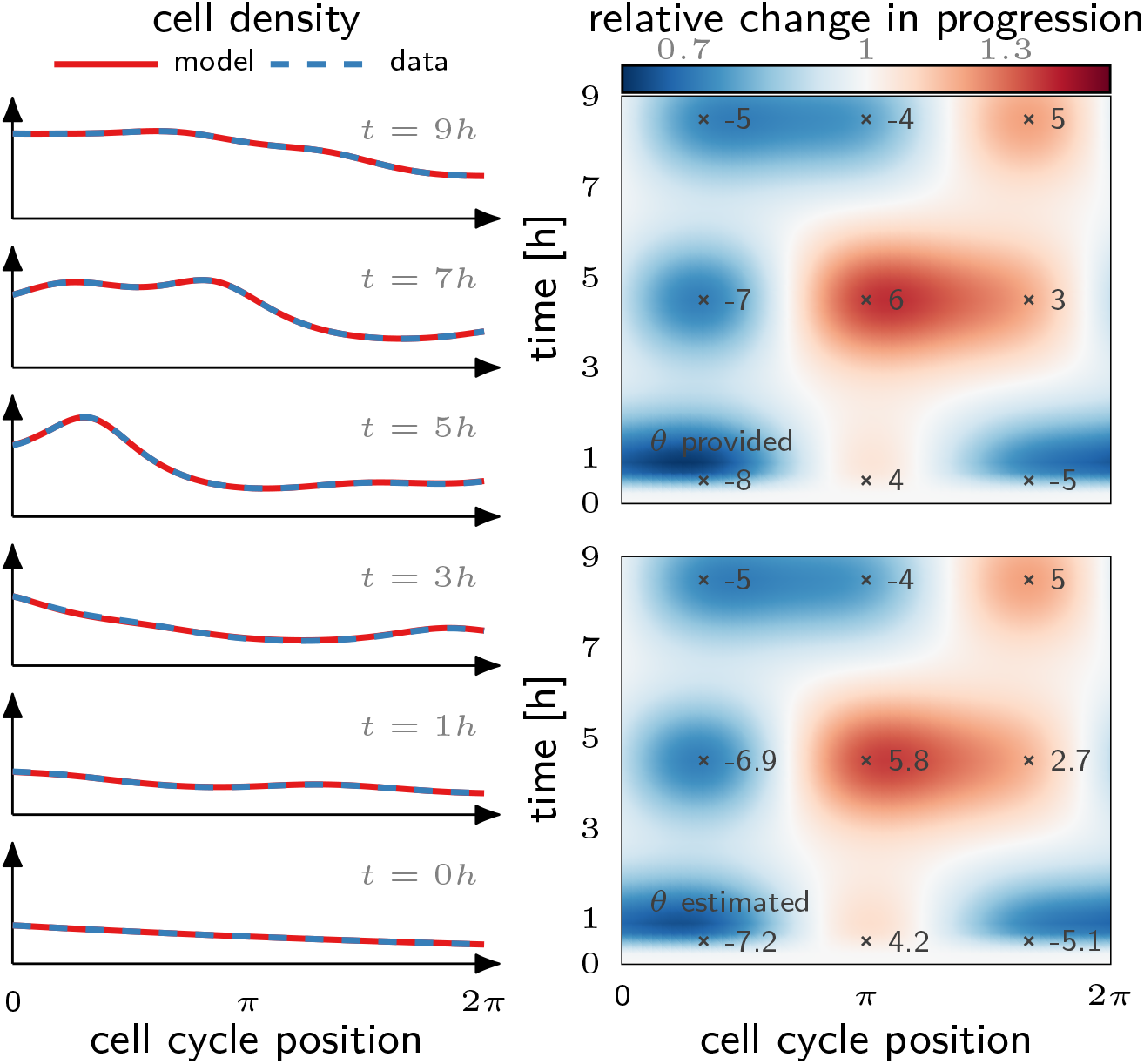
Reconstruction of changes in cell cycle progression from artificial data. After parameter estimation, model predictions (solid, red) perfectly recapitulate the cell densities (dashed, blue) generated from the artificial speed change function (top right). Estimated parameters and the resulting function (bottom right) coincide with the true speed change function and the underlying weights (values shown) for the Gaussian distributions.

### 2.6 Analysis of cell cycle progression

We further tested our method with two proof of concept data sets: (1) control (untreated) and (2) Nocodazole treatment. Nocodazole is an antineoplastic agent that interferes with the polymerization of microtubules, and inhibits the formation of mitotic spindles during M-phase. Therefore we expected that a slow down/arrest of cell cycle progression would occur at the very end of the cell cycle. In contrast, the control experiment was expected to have constant speed change function (*ν*(*t, x*) = 1). Such a constant cell cycle progression would result in the exponential growth of the population with a cell cycle distribution equal to the steady-state Eq. (4).

After preprocessing the experimental data of the control cells as described above, the calculated distributions deviated significantly from the expected steady-state distribution (Figure 4). Accordingly, the estimated speed change function varied strongly depending on time and cell cycle position. In particular, while at later time points, the cell cycle speed in G1 and G2 slowed down, cell cycle progression in the middle of the cell cycle, presumably S-Phase, remained unaltered. We hypothesized that culture conditions during the experiment, such as media exhaustion and increase in cell density, might cause the observed alterations in cell cycle progression.

**Figure 4:**
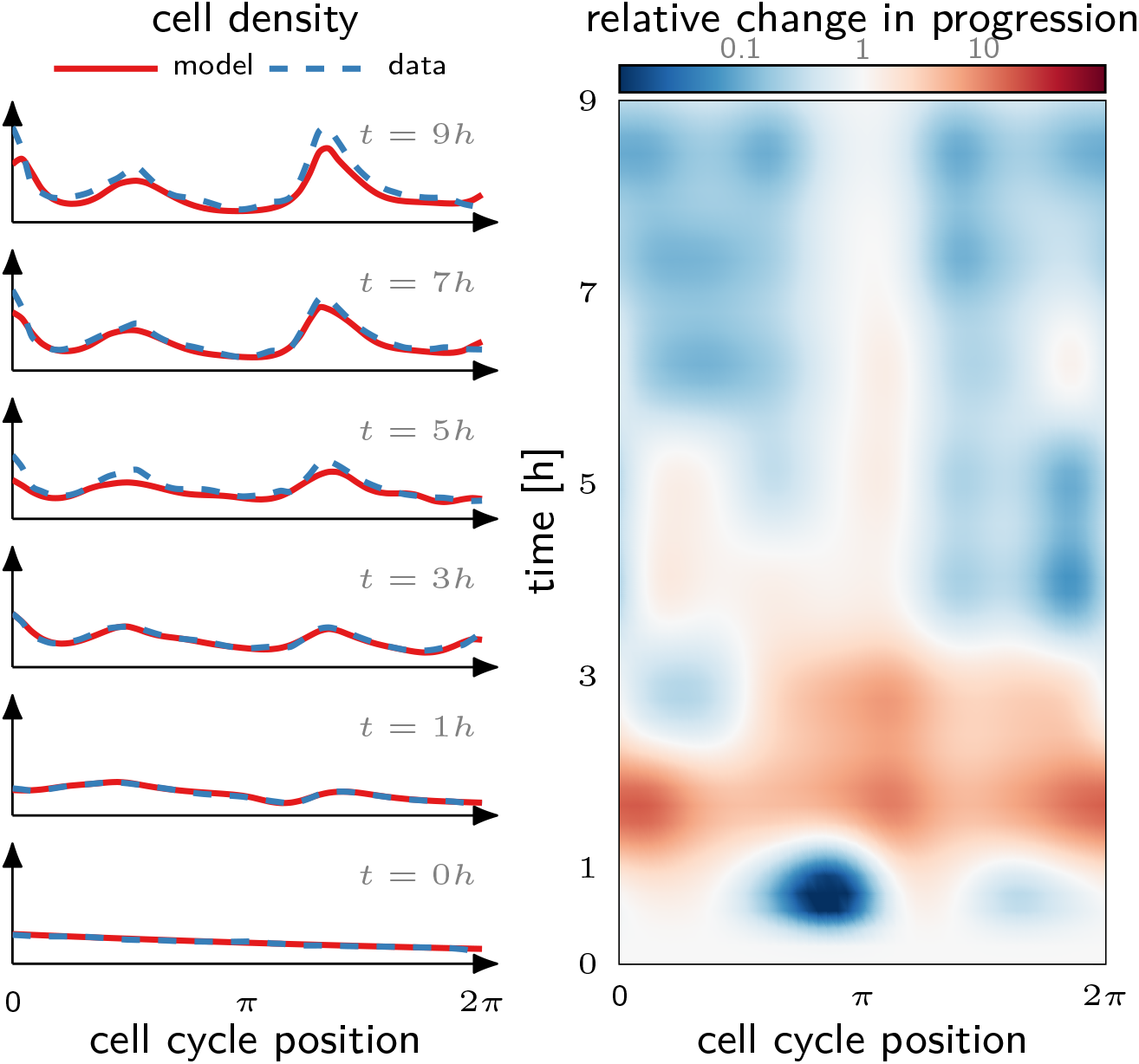
Estimated changes in cell cycle progression in a control experiment with untreated cells. Model prediction (solid, red) recapitulates the experimental data (dashed, blue). Two emerging populations are associated with a slow down of cell cycle progression in G1 and G2, but not S phase.

Cell densities in the experiment with Nocodazole treatment showed a similar pattern with accumulating subpopulations at late G1 and the G2-M transition (Figure 5). However, the population at the G2-M transition increases significantly throughout the experiment. This reduced outflow of cells from G2-M was confirmed by the estimated speed change which showed an almost complete arrest at the end of the cell cycle. This behavior is in line with the established mode of action of Nocodazole. Accordingly, and in contrast to the control experiment, the cell density at the beginning of the cell cycle approached zero, further indicating an inhibition of cell division.

**Figure 5:**
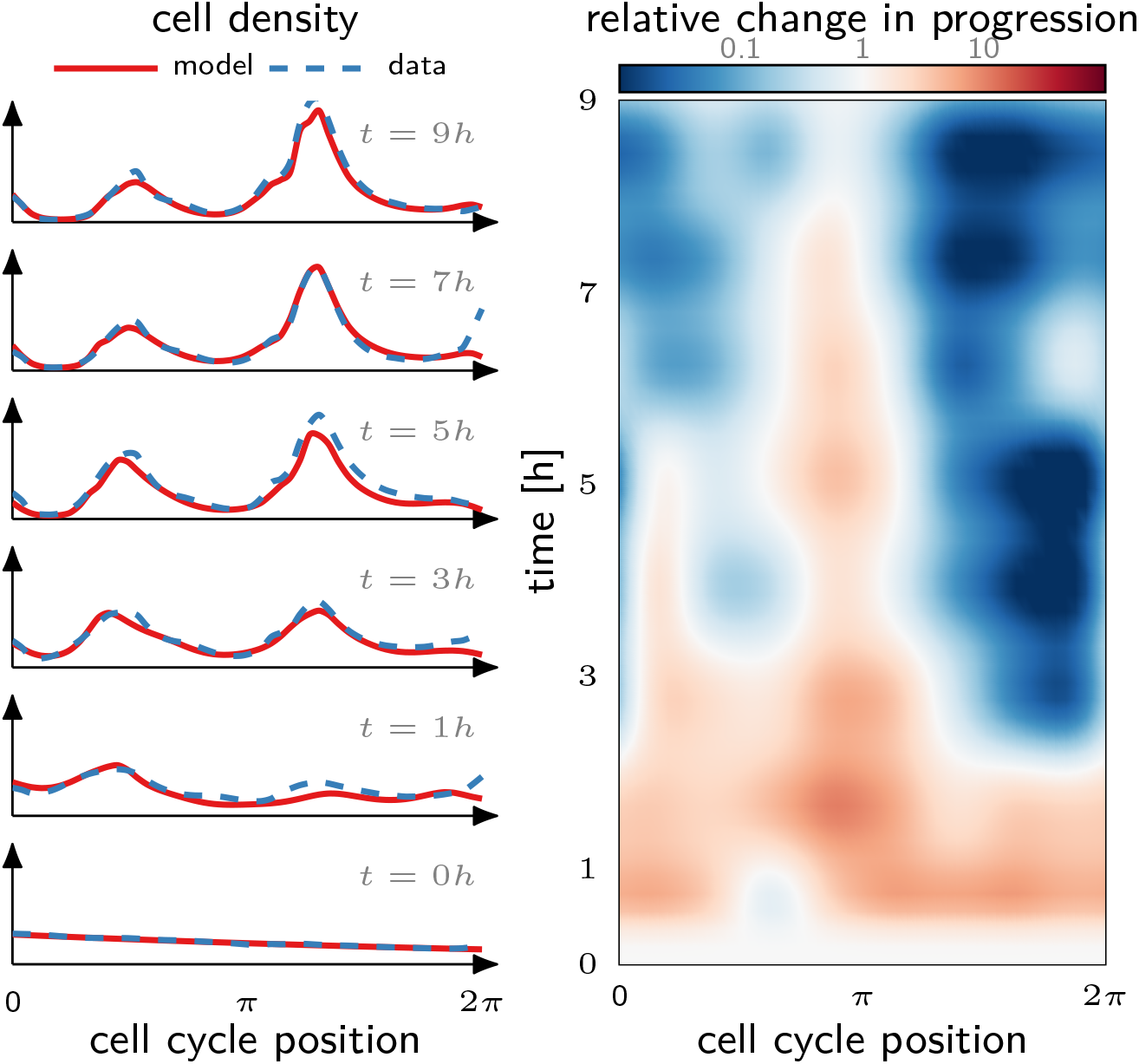
Estimated changes in cell cycle progression a cell population treated with 100*ng/ml* Nocodazole. Model prediction (solid, red) recapitulates the experimental data (dashed, blue). Accumulation of cells at the end of the cell cycle is caused by complete arrest during M-phase which is inline with Nocodazole mode of action.

## Discussion

In summary, we present a computational framework that allows efficient inference of changes in cell cycle progression. To achieve this, we derived the sensitivity system of parameters in the progression function for age-structured type population models. This system allowed the efficient optimization of the objective function, thereby minimizing the difference between experimental observations and model predictions. We showcased the capacity of the method to recover unknown parameters with simulated data. When applied to two experimental data sets, our method uncovered new insights. First, it recognized in the estimated speed change function that Nocodazole induced cell cycle arrest in M-phase. Second, the effects of batch culture cause significant changes in cell cycle progression. Our experimental setup is common for a large class of experiments, and we showed that cell cycle progression is affected by these conditions. Our method can be used to monitor the population for such side effects.

The presented framework serves as a basis and can be extended or modified to broaden its applicability. A mixture model approach, for example, where two or more populations react distinctly to the same stress could be realized by merely weighting the contribution of these populations to the objective function. In this way, one could overcome assumption 3 on the homogeneity of the population’s response. While the descriptive speed change function does not provide mechanistic insights about the cellular response, we envision substituting the speed change function with a mechanistic cell cycle ODE. Such method would require a mapping from the ODE state space to a cell cycle position. The concept of isochrones for nonlinear oscillators provides the theoretical foundation for the existence of such mappings (Winfree 1974; Karsten Kuritz, Halter, et al. 2018). By observing changing dynamics over time one may furthermore trace the rewiring of cellular regulatory networks in response to treatments at different cellular contexts, for example, cell cycle stages.

## Materials and Methods

K562 cells expressing mCherry fused to Cdt1 and Geminin(1-110) fused to EGFP were grown overnight in Dulbecco’s Modified Eagle Medium supplemented with 5% fetal calf serum in 96-well plates (Corning) with an initial seeding density of 5E4 cells/mL. 100 *μL* samples were collected every 2 hr, and cells were stained with Hoechst 33342 Fluorescent Stain (16 *μ*M) for 20min at 37C. Cells were analysed on a BD LSR2 flow cytometer. DAPI (excitation at 355 nm, emission at 450nm), mCherry (excitation at 561 nm, emission between 610 and 620 nm), EGFP (excitation at 488 nm, emission at 530 nm) and total number of events were recorded for 60 *μ*L of sample.

We measured the relative cell number by taking sample from the well mixed cell culture at each time point. By counting the number of cells with the flow cytometer we obtained the cell density which, converted to relative cell number by taking the culture volume into account.

## Acknowledgments

We thank the Lim Lab for access to their flow cytometer.

